# A method for large scale implantation of 3D microdevice ensembles into brain and soft tissue

**DOI:** 10.1101/2020.03.06.979294

**Authors:** Stefan A. Sigurdsson, Zeyang Yu, Joonhee Lee, Arto Nurmikko

**Affiliations:** School of Engineering, Brown University, Providence, RI 02912 USA; Department of Neurology, Massachusetts General Hospital, Boston, MA 02114 USA; Department of Physics and Astronomy, West Virginia University, Morgantown, WV 26506 USA; Department of Neuroscience, West Virginia University, Morgantown, WV 26506 USA

## Abstract

Wireless networks of implantable electronic sensors and actuators on the microscale (sub-mm) are being explored for monitoring and modulation of physiological activity for medical diagnostics and therapeutic purposes. Beyond the requirement of integrating multiple electronic or chemical functions within small device volumes, a key challenge is the development of high-throughput methods for implantation of large numbers of microdevices into soft tissues with minimal damage. To that end, we have developed a method for high-throughput implantation of ∼100-200 μm size devices which are here simulated by proxy microparticle ensembles. While generally applicable to subdermal tissue, our main focus and experimental testbed is the implantation of microparticles into the brain. The method deploys a scalable delivery tool composed of a 2-dimensional array of polyethylene glycol tipped microneedles which confine the microparticle payloads. Upon dissolution of the bioresorbable polyethylene glycol, the supporting array structure is retrieved and the microparticles remain embedded in the tissue, distributed spatially and geometrically according to the design of the microfabricated delivery tool. We first evaluated the method in an agarose testbed in terms of spatial precision and throughput for up to 1000 passive spherical and planar microparticles acting as proxy devices. We then performed the same evaluations of particles implanted into the rat cortex under acute conditions and assessed the tissue injury produced by our method of implantation under chronic conditions.

## 1. Introduction

One vision of next generation electrical and chemical *in vivo* biointerfaces is the idea of an ensemble of implanted active microscale devices forming a smart body-internal wireless network for sensing and stimulation of the underlying biological circuits. One actively pursued concept for developing large-scale neural interfaces, including efforts in our laboratories, envisions ensembles of wireless autonomous microdevices spatially distributed in the brain^1-4^.

Concurrent with active microdevice development, we are presented with the question as to how these devices could be implanted in a safe, scalable, high-throughput manner. As of today there exist no reported methods to our knowledge for large scale implantation of microscale devices. In this paper we report of the development of means to implant large numbers of ∼100-200 μm scale devices. The method of implantation employs an array of microneedles carrying ensembles of devices as payload, with a bioresorbable polymer, polyethylene glycol (PEG), enabling their release upon implantation into tissue. The devices are constrained within PEG constructs which are shape-optimized for penetration and mounted onto the tips of a supporting array structure enabling parallel implantation of potentially large numbers of microdevices.

Throughout development of the method we have used microparticles in the form of spherical polymer microspheres and passive planar silicon chiplets as proxies for real microdevices. Microfabricated arrays, each carrying up to a thousand such microparticles, have been tested in an agarose model to assess the spatial distribution and precision of microparticle delivery in full 3 dimensions. We have performed experiments in rats in which we implanted ensembles of microparticles into the cortex for assessment of our method *in vivo* in terms of both spatial precision, as well as the resulting tissue injury. Here we report the results of these experiments, and discuss the critical issues involved in enabling effective large scale implantation while minimizing the associated cortical injury.

## 2. Background

In early phases of the research, we investigated different approaches to implantation of microparticles as candidates for further development. As a baseline, we first performed a number of experiments exploring implantation by injection from a hypodermic needle. We noted how various groups had described methods of syringe implantation for delivery of probes and other materials into brain tissue^5-8^. We utilized a gauge 27 needle (410 μm outer diameter, 210 μm inner diameter) as the microparticle carrier, with its piston running the length of the needle for ejection of single ∼100-200 μm size particles that were individually loaded into the needle tip. We implanted particles (∼200 μm diameter) to 25 locations one by one in the rat cortex to a depth of 1.4 mm in a chronic experiment but found that all the particles had returned to the cortical surface after a week-long period *in vivo*. The finding that particles could not be properly secured in the tissue, likely finding a return pathway caused by injury from the necessarily large outer diameter of the syringe needle, lead us to reject this method. Ideally, the diameter of the implantation track should match the diameter of the device. Equally important for us, the method should be readily scalable to large numbers of implantation sites (thousands) throughout the target tissue.

We then built a small benchtop apparatus resembling a “gene-gun”^9,10^ that allowed microscale devices to be accelerated through a pressurized nozzle up to a speed exceeding 100 m/s. This enabled ballistic implantation of ∼100-200 μm size microparticles into the cortex in a rat model to a depth of ∼1 mm, but the excessive collateral tissue injury we observed in this case led us to reject this method as well.

We then arrived at the idea of incorporating PEG as a key material component into a custom designed and microfabricated implantation tool. In a related context, bioresorbable, naturally body-dissolving PEG has been used to stiffen flexible microelectrodes for improved penetration into brain tissue^11-17^, for implantation of a single, or a handful of intracortical electrodes at once. In our case, PEG acts as a carrier for release of a payload of microdevices as opposed to flexible microelectrodes, and the method is compatible with high-throughput implantation into soft tissues. Below, we describe the implantation process in more detail, as well as the process of microfabrication of the implantation tool.

## 3. Results

### 3.1. Implantation mechanics

Figure 1 provides an overview of the implantation mechanics. Each microparticle, a proxy for an eventual active device, is encapsulated within a needle-shaped PEG construct which constrains the microparticle to the tip of a supporting silicon shank. This configuration is replicated across each of the shanks of the supporting array structure (only one shank is shown in figure 1 for clarity). Once the structure is positioned, it is driven into the tissue by a pneumatic inserter which brings the microparticles to their target locations in a fraction of a second. Upon immersion into the aqueous environment the PEG constructs dissolve within one minute. This leaves the microparticles embedded in the tissue and allows the supporting silicon shank array to be retrieved. The method is designed to be scalable to potentially hundreds of lateral locations across the target surface. In our studies, each PEG construct contained up to 10 vertically separated microparticles yielding a 3D distribution of particles within the implanted tissue. Specifically, for an array of 100 shanks, up to 1000 microparticles could be implanted in a single insertion step as shown below. Note that, for illustrative purposes, the process is displayed in video form in supplementary video 1 for a small 3D distribution of microspheres implanted slowly into agarose gel.

**Figure 1.**
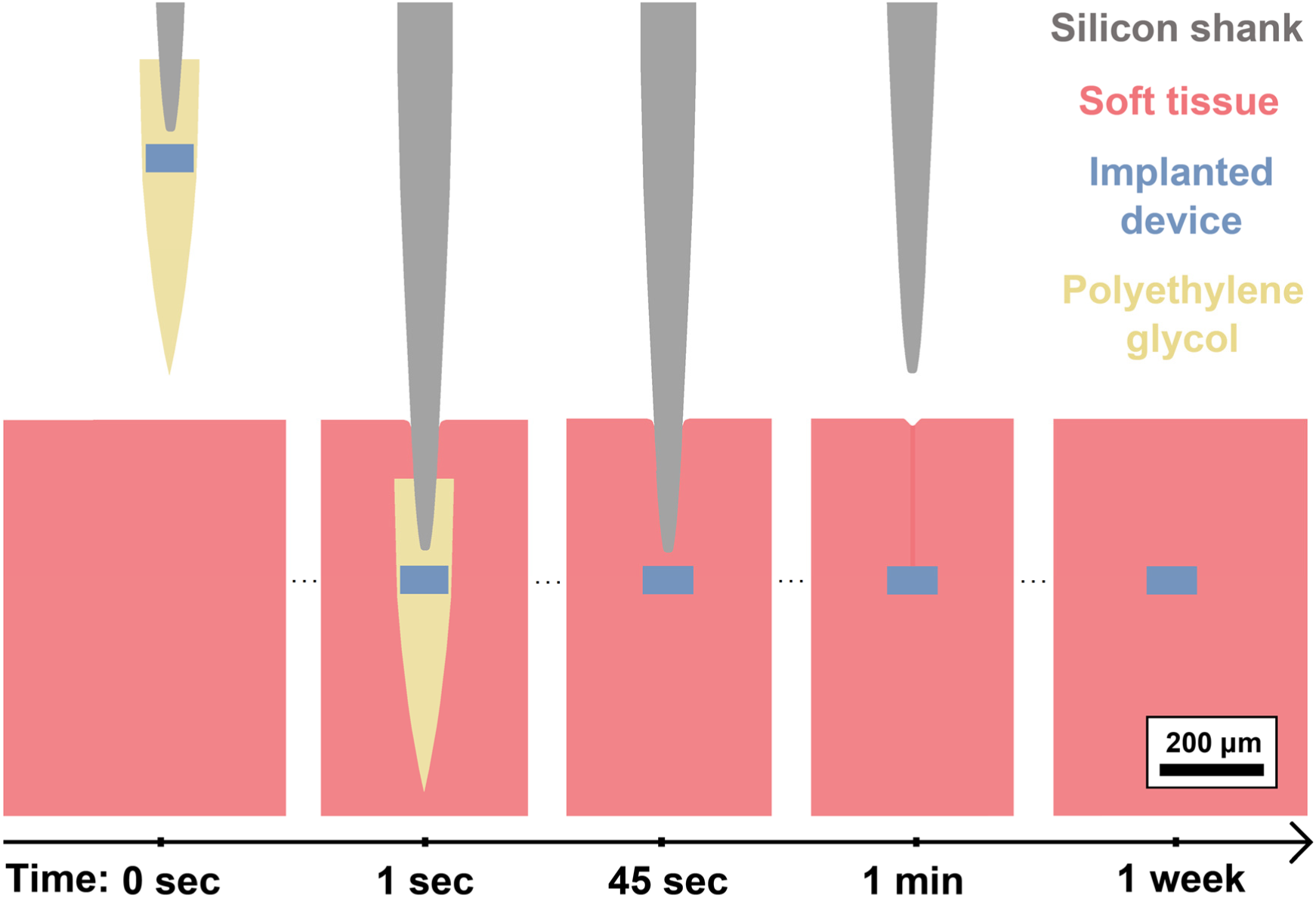
Illustration showing the implantation mechanics. The polyethylene glycol (PEG) mediated process by which microparticles (substitute microdevices) are implanted into tissue is depicted as a time-course. Here, one particle is seen encased within a needle-shaped PEG construct. Upon immersion into tissue (aqueous environment) the PEG dissolves within the course of one minute. The particle is then left embedded in the tissue and the supporting structure is retrieved. While depicted here for a single particle only, each PEG needle may carry several particles stacked vertically, and the process is carried out in an array form for parallel implantation.

### 3.2. Microfabrication of the implantation tool

The microfabrication process by which ensembles of microparticles are loaded into the supporting microarray frame is shown schematically in figure 2. The PEG constructs derive their needle-like shape from a positive stamp used for molding of molten PEG. An important step in the process is the formation of this stamp which we accomplished by two distinct processes. One process choice which we focus on here is silicon based, involving mechanical dicing and wet etching of bulk single crystal silicon to produce the tapered needle shape. In short, a 2.25 mm thick silicon wafer was diced into several shank arrays, with each shank measuring 225 μm by width and 1500 μm by height at 500 μm separation between shanks (center-to-center). The arrays were then wet etched in a 1:19 ratio solution of 50% hydrofluoric acid and 70% nitric acid. The etching time was varied across the arrays to produce an assortment of subtly different shapes. Various groups describe the details of this process for production of microelectrode arrays, such as for intracortical neural recording^18-21^.

**Figure 2.**
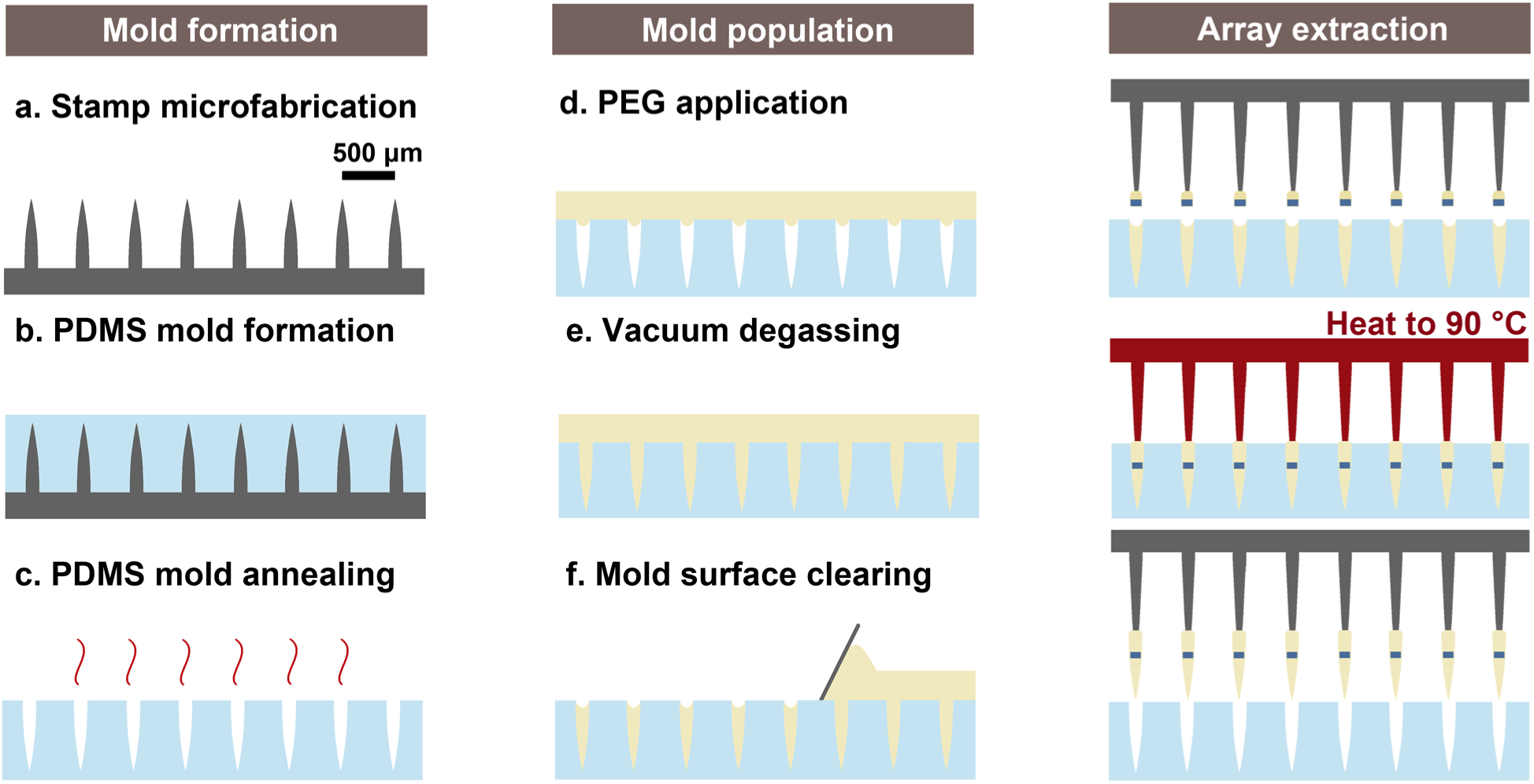
Illustration showing the microfabrication process. The process by which implantation tools are fabricated proceeds as follows. First, an inverse poly-dimethyl-sulfide mold of a microfabricated stamp is used to shape polyethylene glycol (PEG) into sharp needles. The mold is immersed in molten PEG at 90 °C in vacuum for degassing to allow the PEG to conform to the shape of the mold. Once the mold surface is cleared, microparticles are either placed directly into the molten PEG constructs or mounted separately onto the supporting array as displayed here. Finally, the needle-shaped PEG constructs are retrieved from the poly-dimethyl-sulfide mold and mounted onto the supporting array structure by micromanipulators.

The second process, to be described more fully in a later publication, takes advantage of 3D nanoprinting technology by which stamps were formed through the layer-by-layer polymerization of photoresist (Photonic Professional GT system, Nanoscribe GmbH, Karlsruhe, Germany). We produced arrays of needles using the IP-S photoresist (Nanoscribe IP-S, Nanoscribe GmbH) at 1 μm process resolution, with each needle measuring 125 μm by width and 550 μm by height at 500 μm separation between needles (center-to-center). The details of the process resemble those reported by Faraji Rad et al.^22^ This method of stamp microfabrication was explored as a more accessible and customizable option for custom stamp fabrication over the silicon based process. The silicon microfabrication process had the advantage in terms of throughput however as the nanoprinting process, by currently available moderate throughput and moderate cost equipment, would take several hours to produce even a relatively small stamp form factor (6×6 array, short needles).

Once the appropriate stamp was obtained, it was used for production of reusable poly-dimethyl-sulfide (PDMS) molds. PDMS (Sylgard 184, Dow Corning, Michigan, USA) was mixed at 1:7 ratio of hardener to elastomer for production of stiff molds. Once degassed, the PDMS was cured at 50 °C for 6 hours to form a mold around the stamp, and each mold then annealed at 120 °C for 24 hours after the stamp was retrieved. At this stage, the molds were immersed in molten PEG and a vacuum oven used for degassing at 90 °C for 10 minutes, allowing the PEG to conform to the shape of the molds. The PEG is rendered molten at ∼65 °C, so an elevated process temperature is necessary. As for the choice of large PEG molecular weight of 35 kDa (CAS 25322-68-3, Sigma Aldrich, Missouri, USA), the reader is referred to discussion by Lecomte et al.^11^

Once the surface of the molds was cleared, microparticle populations were either loaded directly into the molds, or mounted onto the supporting shank array in a separate process by a simple adhesion step as depicted in figure 2. The supporting shank array was then aligned with the PEG populated mold by micromanipulators, and the PEG constructs mounted onto the supporting array by temporarily rendering the PEG molten. While the PEG is in a molten state, the supporting shanks are able to slide into the constructs and attach as the PEG cools down and solidifies. The PEG constructs are then retrieved from the mold by raising the supporting array to complete the microfabrication sequence. Note that in order to consistently retrieve the molded PEG constructs from the molds without breaking at 100% yield, it was crucial that the molds be stiff and thoroughly annealed.

Figure 3 provides a view of the positive stamps produced by the two methods discussed above, as well as the corresponding PEG constructs obtained through molding. The PEG constructs here are mounted onto the tips of supporting arrays, which are themselves produced through a similar process as described for the silicon stamp above. We have performed SEM imaging of the stamps, as well as their PEG counterparts, for assessment of the degree to which the molding process accurately replicates the shape of the original stamp, specifically in terms of the needle tip size and surface roughness. For each sample, a measurement from 30 needles was obtained. In case of the silicon based process, the stamp tip size measured 2.13 ± 0.47 μm (mean ± std), in contrast to 2.37 ± 0.50 μm measured for the corresponding PEG needles. For the 3D nanoprinting process, the stamp tip size measured 1.25 ± 0.25 μm, with the respective PEG needle tips measuring 1.53 ± 0.32 μm. Surface roughness measured under 1 μm in all cases. The shape and surface roughness of penetrating probes have been shown to affect the extent of initial tissue injury caused by implantation^23,24^, so it is important that these parameters translate effectively from microfabricated stamps to the PEG constructs.

**Figure 3.**
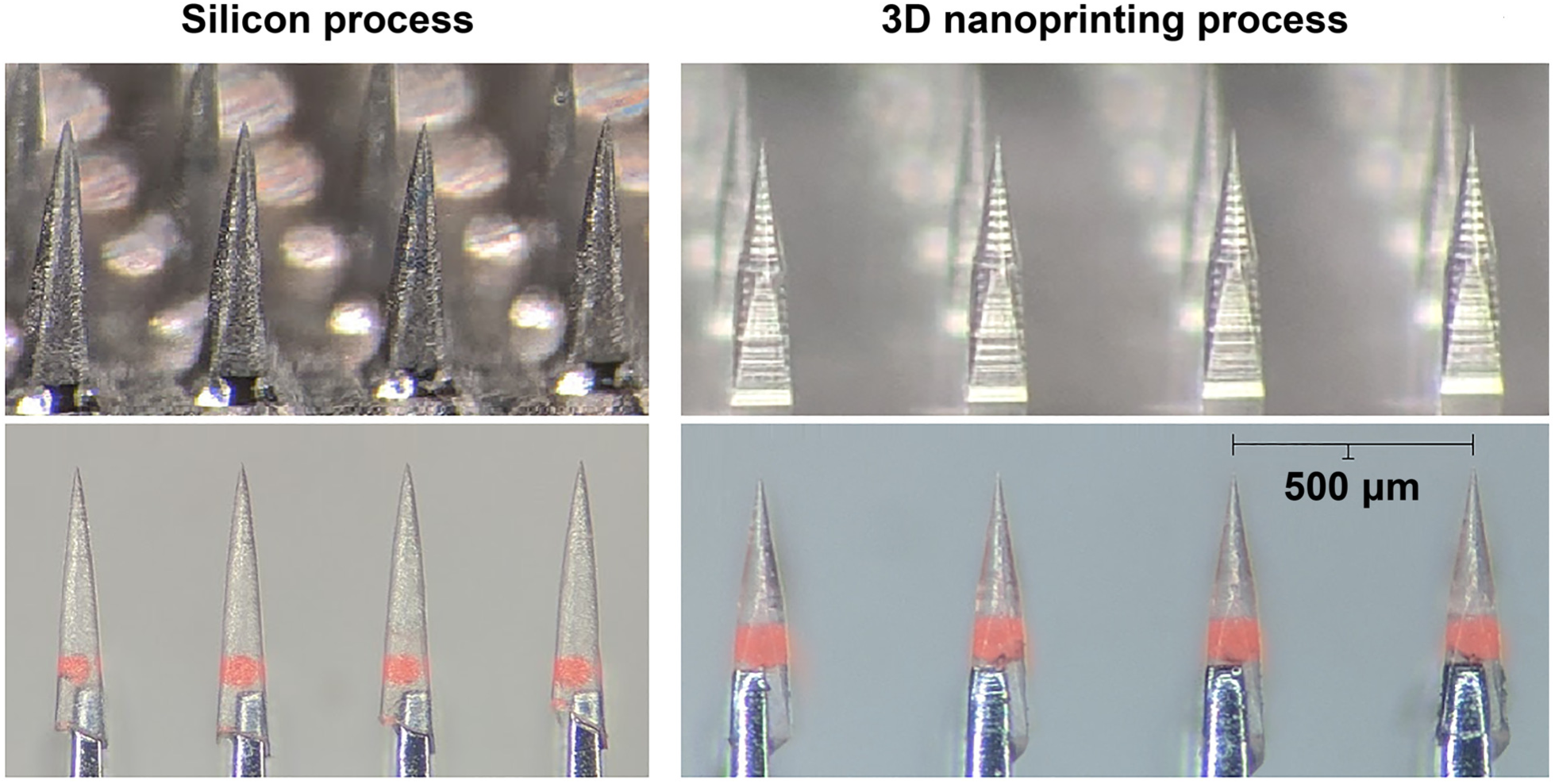
Comparison of stamp replication fidelity. Images of microfabricated stamps along with the corresponding molded polyethylene glycol constructs depicted for the two different stamp fabrication processes (silicon wet etching on the left, 3D nanoprinted structure on the right). The stamps are displayed in the top two panels, while the corresponding polyethylene glycol needles are displayed in the bottom two panels carrying fluorescent microspheres and mounted onto the supporting array structure. Inter-needle separation measures 500 μm in each case.

### 3.3. Spherical and planar microparticles as substitute microscale devices

In lieu of active electronic microdevices or chemically infused microscale payloads, we chose two types of passive proxy particle types to study the effects of different geometries and material choices; (a) spherical polyethylene and borosilicate microspheres of size ∼100 μm were purchased from Cospheric (UVPMS-BR-1.090, HCMS-BSGMS-FMB, Cospheric, CA, USA), and (b) planar silicon chiplets measuring 100×100 μm on the side and 50 μm thick were microfabricated through a combination of photolithography and dry etching. To describe the microfabrication process, a 50 μm thick silicon wafer was first mounted onto a carrier wafer by a thin layer of thermal grease (AOS 340 WC, AOS Thermal Compounds, NJ, USA). An S1813 photoresist mask (Microposit S1813, Shipley Company, MA, USA) was patterned onto the wafer, and the wafer etched through by deep reactive ion etching (Omega LPX Rapier, SPTS Technologies, Newport, United Kingdom). The sectioned microparticles were then released in acetone and cleaned by ultrasonic cleaner (Elmasonic P, Elma Schmidbauer GmbH, Singen, Germany). We chose the particular size range near 100 μm recognizing ongoing efforts to create micro- and optoelectronic microscale active implants for biosensing and actuation. We have also performed experiments using 200 μm size microspheres but as those produced qualitatively similar results as obtained for the smaller sizes, we will not discuss them further. Figure 4 summarizes the microparticle shape and size for the two types in our study.

**Figure 4.**
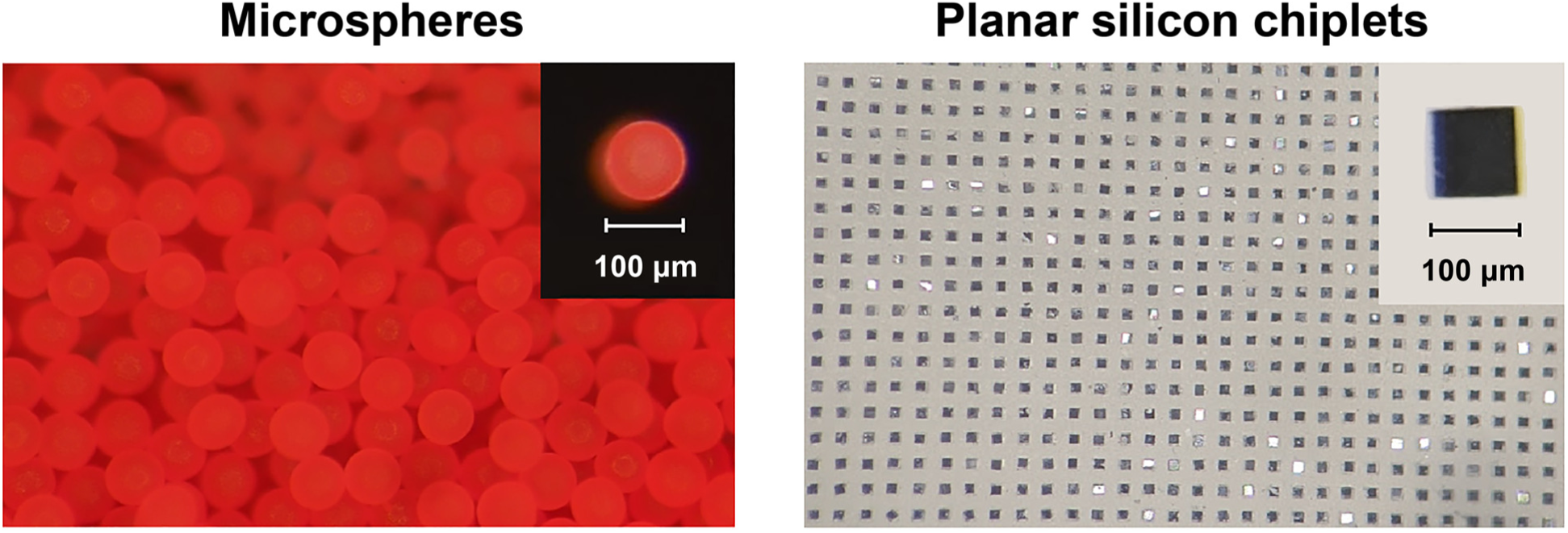
The substitute microdevices used. The two types of substitute microdevices (microparticles) used in our experiments are displayed (fluorescent microspheres on the left, planar silicon chiplets on the right). The two types present different geometries which allow us to study geometry dependent behaviors. The left hand side displays an unordered collection of particles as received from the vendor, while the right hand side displays particles organized by the use of a molded “waffle pack”. Scale factor is shown in the inset for each type of microparticle.

To guide the process of loading and mounting microparticles onto supporting arrays, PDMS molds in the form of a “waffle-pack” were used to organize unsorted microparticles into a regularly spaced grid matching that of the stamps. Figure 4b displays sorted planar microparticles residing in such a mold, whereas figure 4a shows an unordered collection of microspheres as received from the vendor. The molds were formed using a stamp produced by patterning of SU-8 photoresist (SU-8-2100, MicroChem Corporation, MA, USA) into regularly spaced 100×100×100 μm blocks on a silicon wafer. Microparticles were organized using these molds by simple mechanical agitation of particles placed on the mold surface (note that molds must be thoroughly annealed for this purpose). The particles could then be mounted in parallel onto supporting arrays by a small amount of PEG at the supporting shank tips. This allowed for high-throughput placement of both the microspheres and the planar chiplets into the PEG constructs, at a rate of approximately a thousand microparticles arranged for every few minutes of effort.

### 3.4. Unordered implantation of 3D microsphere ensembles into agarose

There are several ways by which groups of microparticles may be placed into the molded PEG constructs during preparation of the implantation tool. If one wishes to implant large numbers of microparticles (≥1000) across a 3-dimensional target volume, several microparticles can be placed within each PEG shank. A quick way of placing the microparticles into the molds is to simply pack the molds with particles to capacity before the application and vacuum degassing of molten PEG. In this way, thousands of particles may be placed into a mold in a matter of minutes, although this approach results in a spatially unordered 3D distribution of microparticles after implantation. The results of this process are depicted in figure 5 in which an array composed here of 10×11=110 PEG constructs was implanted into agarose, delivering a total of ∼600 microspheres without the assistance of an additional supporting structure. Indeed, the use of a supporting array is not crucial to the process, and for some applications foregoing it may be possible depending on physical characteristics of the tissue in question and the desired implantation depth. We have found that the microspheres typically self-align along the vertical axis within each PEG construct, with a small spacing of 5-20 μm between each pair of microparticles as seen in figure 5c. In some cases, surrendering control of the inter-particle spacing and relying on this type of quasi-random self-assembly along the vertical axis may produce acceptable outcomes.

**Figure 5.**
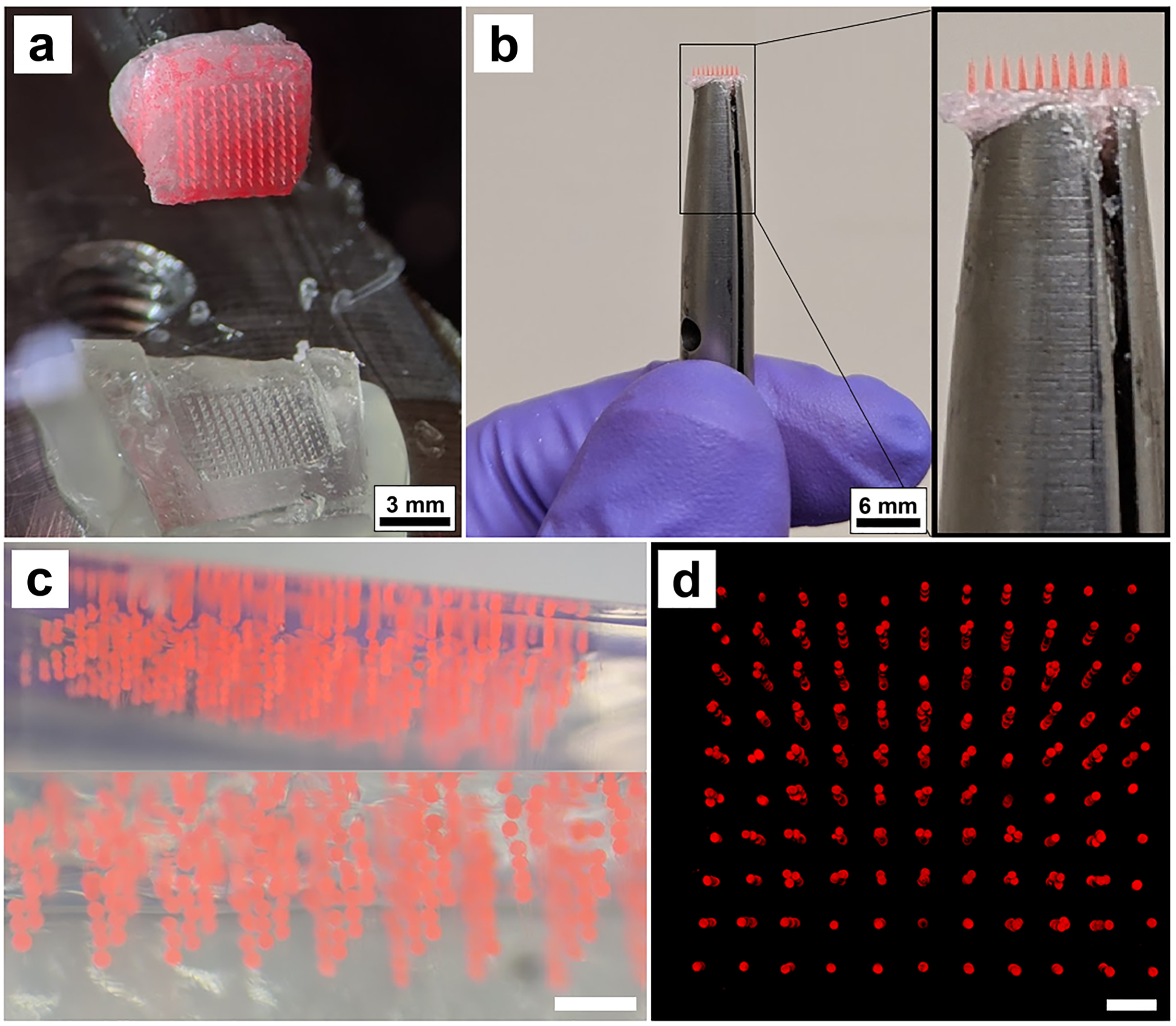
Unordered microparticle implantation. An unordered ensemble of ∼600 microspheres was implanted into agarose. **a** Retrieval of the polyethylene glycol constructs from the mold. Note the omission of a supporting array structure, here replaced by a simple polyethylene glycol base. **b** Implantation tool held by hand for perspective. **c** The ensemble of microspheres implanted into agarose gel which is displayed from a side view. **d** Reconstruction of the microsphere ensemble in agarose by confocal fluorescence microscopy displayed from a top view. Scale bars in (**c**) and (**d**) measure 500 μm.

Following implantation, the agarose gel was imaged by confocal fluorescence microscopy for reconstruction of the microsphere ensemble within the gel. The 3D reconstruction is displayed in figure 5d and is available in video form in supplementary video 2. While vertical self-alignment is prevalent, some aggregation is seen towards the top of the vertical implantation columns in figure 5d. The needles of the stamp used for generation of the PEG constructs in this experiment tapered to ∼200 μm width at their base, thus leaving some room for particles to be displaced from perfect vertical alignment near the base of the PEG constructs. We measured this horizontal displacement for each particle (from their respective implantation columns) of the ensemble and found that the average displacement (center-to-center) measured 28 μm, with 90% of particles displaced less than 55 μm. We believe this displacement, as well as the aggregation effect, can be significantly reduced with the use of a more optimized stamp shape. Nonetheless, these experiments demonstrate that the spatial distribution of microparticles is not significantly altered by the process of PEG dissolution within the agarose gel.

### 3.5. Ordered implantation of planar silicon chiplet ensembles into agarose

Controlling not only the inter-particle spacing, but also the individual microparticle orientation may be important for active implants (e.g. antenna orientation for energy harvesting by inductive coupling^1^). Placement of microparticles in an ordered manner with specific orientation into the PEG constructs introduces additional complexity, especially in terms of controlling inter-microparticle spacing for multi-layered (3D) implant populations. Using the “waffle-pack” described in section 3.3, microparticles in the form of planar chiplets may be oriented and spaced appropriately in large numbers with minimal effort for high throughput placement into the PEG constructs. For a single 2D layer of chiplets the process is relatively straightforward. If several chiplets are to be stacked in depth within the same PEG construct, however, the process must be repeated several times with additional embedded spacer layers of correct thickness alternating with the chiplets. One convenient spacer material is Silk Fibroin. Microfabricated slabs of bioresorbable Silk Fibroin can endure the repeated heating of PEG to 90 °C, while dissolving over time once implanted into tissue^25-27^. In this manner, ordered implantation of large numbers of microparticles can be achieved, although the effort involved in this case is proportional to the number of layers desired. As displayed in figure 6, the orientation of microparticles, as well as inter-microparticle spacing, can be retained through the process of implantation into agarose. As an alternative to the layer-by-layer process, pre-processing of microparticles into columnar constructs for single-step placement into the molds could greatly reduce the effort involved. This process would need to be uniquely tailored to each kind of microparticle, or microscale device, however.

**Figure 6.**
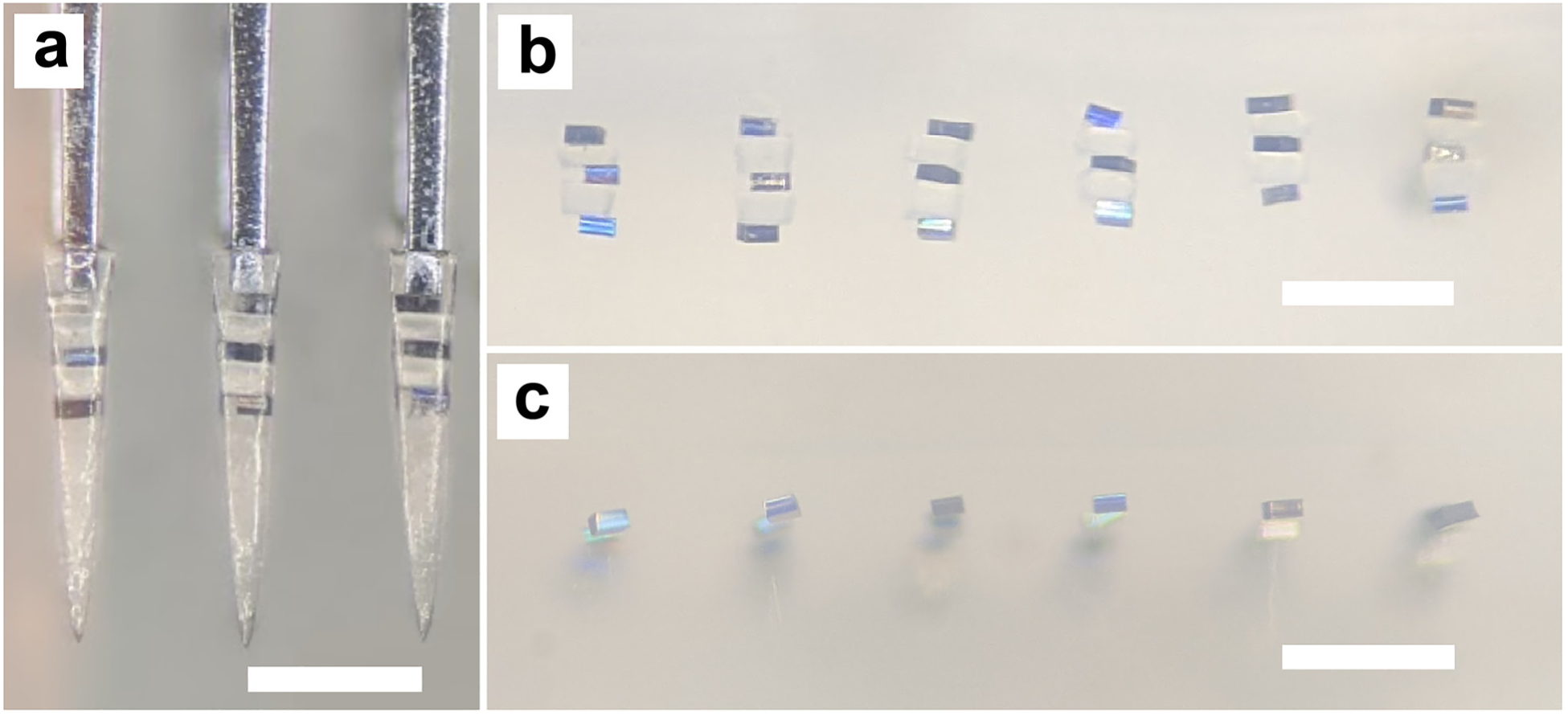
Ordered microparticle implantation. Ordered ensembles of planar silicon chiplets were implanted into agarose. **a** Multi-layered ensemble of regularly spaced chiplets encased within PEG needles prior to implantation. **b** Side-view of the same ensemble implanted into agarose. **c** Side-view of an implanted single-layer ensemble in agarose. Note the extent to which particle orientation and spacing is preserved on implantation. Scale bars measure 500 μm.

### 3.6. Acute in vivo experiments: Assessment of placement accuracy

As the spatial distribution and orientation of microparticles was adequately conserved on implantation into agarose, the question followed as to whether the same applies to *in vivo* conditions, given the multiple biophysical forces which could affect the outcomes in living (brain) tissue. An acute rat model is a useful testbed even if the size of the animals’ head/brain limits the overall implantable payload. We implanted a single-layer (2D) ensemble of 36 planar silicon chiplets into the rat cortex in an acute experiment. Separately, we also implanted a multi-layered 3D ensemble of ∼260 glass microspheres in a 6×6 array form factor into the rat cortex to study implantation of 3D particle distributions (number of particles constrained by the head size of the rat model). The implantation process is depicted in figure 7a, with the inserter and the implantation tool in view above the cortex. We found that penetration into the cortex is consistent, with little or no dimpling of the cortical surface observed. In terms of overt tissue damage, we found that it was comparable to that of implantation of conventional microelectrode arrays immediately following implantation. Minimizing acute tissue damage is especially important as we have found earlier how e.g. larger diameter hypodermic needles create damage tracks in brain tissue which act as channels of particle ejection. We note that in our chronic experiments we did not observed particle ejection using the present method, in contrast to the hypodermic needle injection method.

**Figure 7.**
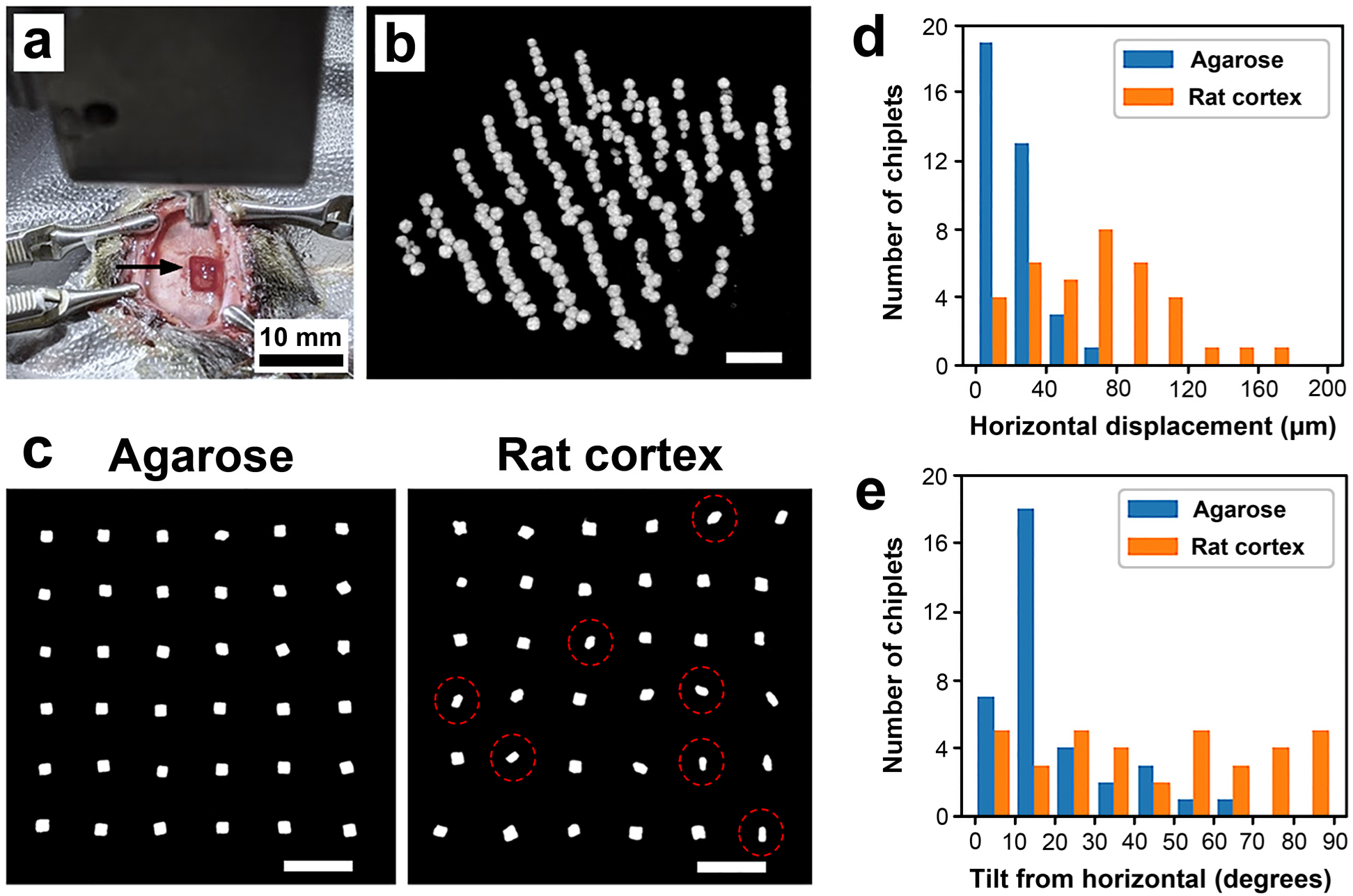
Implantation precision assessed *in vivo*. A single-layer ensemble of planar silicon chiplets and a multi-layered ensemble of glass microspheres were acutely implanted into rat cortex and imaged by micro-CT scanner. **a** The experimental setup is displayed with the inserter and the implantation tool above the target cortical location before implantation. Arrow points to the exposed craniotomy. **b** Reconstruction of an intracortical 6×6 3D distribution of ∼260 glass microspheres is displayed, viewed from an angle. **c** Reconstructions of two 6×6 distributions of planar silicon chiplets are displayed from a top view, one implanted into agarose and the other into rat cortex. Some chiplets that have rotated towards 90 degree tilt are circled in red. **d-e** Outcomes of the experiments in (**c**) were quantified. **d** Histogram showing the horizontal displacement of particles from their respective vertical implantation columns (center-to-center) in agarose and in rat cortex. **e** Histogram showing the deviation of particles in terms of tilt in orientation from the initial horizontal orientation in agarose and in rat cortex. Note the near uniform distribution of particle orientations in the rat cortex. Scale bars in (**b**) and (**c**) measure 500 μm.

The micro-CT scans obtained from the acute *in vivo* experiments involving both spherical (glass) microspheres and planar silicon chiplets were used to form 3D reconstructions of the particle ensembles (see figure 7b-c), which were analyzed similar to that described in section 3.4 above. While the microspheres might be expected to show minimal rotation due to their isotropic geometry, the chiplets possess large shape anisotropy whereby reorientation (tilt) might occur. Information was first gathered about the horizontal (in-plane) deviation of chiplets from their respective vertical implantation columns (center-to-center). We then focused on the deviation of chiplets in terms of tilt of orientation from the initial horizontal plane. The results are displayed in figure 7d-e. Here, data from identical agarose experiments is overlaid to give a sense of the difference between the two experimental conditions. We found that chiplet positioning in the horizontal plane was more robust in agarose (mean 21 μm deviation) compared to *in vivo* (mean 67 μm deviation), perhaps largely due to swelling of the tissue, although both conditions produced satisfactory results for our needs.

As for deviations in chiplet orientation i.e. tilt, the noticeable distribution of tilt angles *in vivo* compared to that in agarose shows how biophysical forces in the living cortex could drive an initial (near) zero-tilt angle distribution towards a 90 degree tilt after implantation. While the driving forces at play in tilting chiplets by 90 degrees are unknown to us, there could be both intrinsic structural reasons (here cortical columns) and extrinsic forces (residue of implantation damage) which favor such an “edgewise” or vertical realignment of the chiplets. For active microelectronic devices such reorientation must be taken into consideration when designing orientation sensitive features such as antenna structures for energy harvesting. A major difference between the brain tissue environment and the agarose is that the tissue is dynamic. The tissue pulses with every heartbeat, expanding and contracting in a way that may influence the orientation of implanted devices. Investigation into the specific cause of the observed dynamical re-orientation is the subject of current exploration which aims to identify strategies by which device orientation may be preserved or otherwise controlled if so needed.

### 3.7. Chronic in vivo experiments: Assessment of cortical injury

For assessment of the cortical injury produced by our system we implanted unordered 3D microparticle ensembles into the cortex in 4 rats to a cortical depth of 800-1400 μm (by use of supporting arrays). Each ensemble totaled between 100-150 fluorescent polymer microspheres in a 5×6 form factor. At 4 weeks post-implantation the animals were perfused and their brains dissected. The brains were consecutively cryosectioned in the horizontal direction at 20 μm slice thickness, and every third slice then stained with β3-Tubulin for labeling of neurons, as well as DAPI for labeling of cell nuclei. The slices were then imaged by fluorescence confocal microscopy and the neuronal and nuclei presence around the implanted microparticles characterized. In the vast majority of cases the sliced fluorescent polymer microspheres were ejected from the tissue slices in the gentle treatment steps prior to imaging (see figure 8a-b), so for identification of particle sites in the cortical slices we looked for the presence of voids larger than 80 μm in diameter (microspheres measured 90-110 μm in size). Voids of this size were not found in the healthy hemispheres of the subjects, while hundreds of such spherical voids were found across the affected hemispheres in a rough 5×6 configuration. For any voids meeting the size criterion we imaged and then calculated the normalized β3-Tubulin and DAPI fluorescence intensities (per pixel) as a function of distance (1 μm bins) from the edge of the microparticle void. Normalization was done with respect to the fluorescence intensity measured at 160-180 μm distance from the microparticle voids. No voids identified measured larger than 110 μm, which is consistent with the size range of the implanted microspheres. Aggregate results are displayed in figure 8c for 30 microparticle voids across 4 rats imaged under standardized conditions. As shown, we did not find a significant drop in β3-Tubulin intensity in proximity to the microparticles, indicating that the proximal neuronal population was not significantly diminished by our implantation method, the PEG, or the chronic presence of the microparticles themselves. We did, however, find an increase in DAPI intensity near the microparticles, indicating the formation of an encapsulating layer of tissue extending 15-20 μm from the microparticle edge. As the neuronal decline, as well as the spatial extent of the encapsulating tissue, was well within normal limits found for other intracortical devices^28,29^ (see e.g. neuronal decline described by Biran et al.^28^), detailed immunological characterization of the encapsulating tissue was not deemed a priority and is left as the subject of future experimentation.

**Figure 8.**
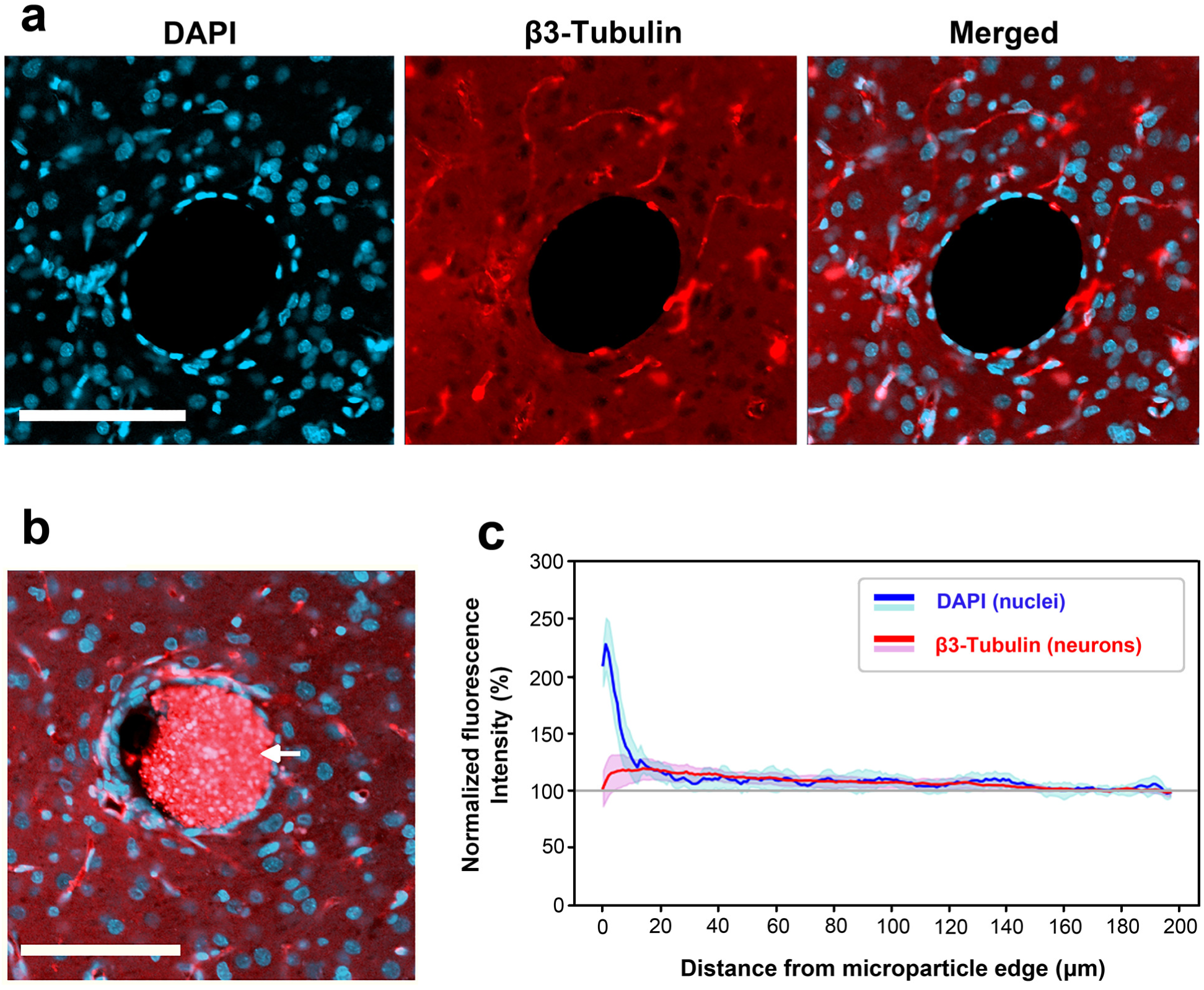
Assessment of chronic tissue injury. Characterization of the tissue reaction to microparticles implanted for 4 weeks in the rat cortex. **a** Representative view of a microparticle void and the surrounding tissue with cell nuclei labeled blue (DAPI) and neurons labeled red (β3-Tubulin). **b** Example of a tissue section with a sliced microparticle still present (arrow). **c** Normalized DAPI and β3-Tubulin fluorescence pixel intensities are plotted as a function of distance (1 um bins) from the microparticle edge. Each trace is the average of data collected from the vicinities of n=30 microparticles across 4 rats. Shaded areas represent one standard deviation. Scale bars measure 100 um.

## 4. Discussion

Development of systems of implantable microscale devices for applications in the body is the subject of ongoing research. While various groups have published efforts towards miniaturization of active devices, a question that remains unanswered is how these kinds of devices would be implanted into tissue in large numbers in a high throughput manner. The method we have described here is applicable to a variety of device geometries and system configurations for high-throughput implantation into the nervous system, as well as generally into other soft tissues. As researchers look for methods to enable implantation of their devices, the method we have developed could be used for production of appropriate implantation tools. With those tools provided, the only components requiring operation at the hands of the researcher would be the micromanipulator mounted pneumatic inserter. For specific custom applications, researchers could opt to produce customized stamps for mold formation with minimal effort by use of a 3D nanoprinter or equivalent (see section 3.2).

Our method has proven reliable in delivering ensembles of microparticles to precise locations in the cerebral cortex, a particularly challenging bio-environment. In terms of tissue injury, we found no signs of neuronal loss around chronically implanted microparticles 4 weeks post-implantation. We did find signs of encapsulating tissue in the same experiments, although extending only 15-20 μm from the surface of the microparticles. While a detailed analysis of the encapsulating tissue was not performed, our results indicate that implantation of microparticles by our system produces tissue injury comparable to that of conventional intracortical devices. While we did assess the spatial precision of our system under acute conditions in a rat model, one question that remains unexplored is whether, and to what extent, this precision carries over to chronic conditions and to primates. In our chronic experiments we found that the implanted particles remained roughly in their original relative configuration (5×6 form factor, 600 um vertical spread), but a detailed assessment of exact particle drift and of change in particle orientation over time could not be made. This, in addition to the additional challenge of safe *in vivo* extraction of implanted microparticles, is subject of current work aimed at exploring the full device lifecycle in chronically implanted animals.

We note that while we have focused on implantation of microscale devices, implantation of arrays of flexible electrodes (or similar physiological probes) is in principle also supported by our method. Further, by way of comparing another advanced implantation approach for embedding microprobes into brain cortex, we note the system being developed by Neuralink^30^. The reported rate of implantation for that system amounts to 6 separate implantation sites across the cortex per minute, although they implant only a single probe at a time. By tailoring our approach, implantation tools of 100 shanks each could conceivably be implanted in a similar manner for a throughput exceeding that reported so far by two orders of magnitude.

## 5. Materials and methods

### 5.1. Assessment of implantation methods

The precision of our implantation methods was assessed first by study of ensembles of microparticles implanted into agarose brain phantom (0.6% w/w in de-ionized water), and then *in vivo* rat cortex. In the case of the agarose testbed (which mimics the mechanical features of the brain^31^) implanted fluorescent microspheres were imaged by confocal fluorescence microscopy for evaluation (FV3000, Olympus, Tokyo, Japan). As for particles implanted into rat cortex, imaging was performed by micro-CT scanner (microCT 40, Scanco Medical, PA, USA) in order to overcome the optical opacity of brain tissue. Both imaging methods allowed for 3D reconstruction of implanted microparticle ensembles at a resolution of a few micrometers. The reconstructions were then used to calculate deviations of the implants from their ideal configurations.

### 5.2. Surgical methods

Evaluation of our implantation methods *in vivo* was carried out as follows. Male Long Evans rats (8 weeks old, obtained from Charles River Labs, MA, USA) were anesthetized by isoflurane (1-3% concentration in oxygen) and mounted onto a stereotaxic apparatus. A craniotomy measuring 4×6mm was performed over one hemisphere between the lambda and the bregma, with the dura mater subsequently removed by sharp tweezers. The implantation tool was then secured over the exposed craniotomy, and a solenoid actuator (inserter) used to drive the array of microparticles into the cortex at high velocity (∼5-10 m/s). Note that high-speed implantation of penetrating arrays has been reported to reduce the dimpling of the cortex on implantation as well as the resulting tissue injury^32-34^. Once the array carrying the microparticles was extended into the tissue, five minutes were allowed to pass to permit diffusion of the dissolved PEG and allow the tissue to conform to the particles. The array was then slowly retracted, and the particles left embedded in the tissue. Animal care and experiments were performed in accordance with the National Institutes of Health guidelines and approved by the Brown University Institutional Animal Care and Use Committee (protocol #1704000267).

### 5.3. Immunohistology

For histological analysis, rats were terminally anesthetized with pentobarbital (intraperitoneal injection). Transcardial perfusion was performed with 0.1 M phosphate buffered saline (PBS), followed by 4% paraformaldehyde in 0.1 M phosphate buffer (PB). The brain was extracted and fixed in 4% paraformaldehyde in PB overnight, and then cryoprotected in 30% sucrose solution (until the brain sunk in the solution). The brain was then frozen by dry ice and sectioned into 20 um thick horizontal slices using a microtome. Brain slices were then washed in PB (twice, 5 min each) and PBS (twice, 5 min each) and left for 1 hour in block solution containing 10% goat serum (ab7481, Abcam, Cambridge, United Kingdom), 0.25% Triton-X 100 (Sigma Aldrich), and 0.1% Tween (Sigma Aldrich) in PBS. The slices were then stained overnight at 4 °C in block solution containing beta III tubulin (β3-Tubulin) antibodies for labeling of neurons (1:100 ab201740 Anti-beta III Tubulin antibodies conjugated with Alexa Fluor^®^ 594, Abcam). The slices were then washed in PBS (twice, 5 min each) and PB (twice, 5 min each) and mounted onto microscope slides with diamidino-2-phenylindole (DAPI) mounting medium for fluorescence. Finally, the slides were imaged by confocal fluorescence microscopy for evaluation (FV3000, Olympus).

## Acknowledgements

This research was supported in part by the DARPA program Neural Engineering Systems Design (NESD) (N666001-17-C-4013) and a private gift to Neuroengineering research at Brown University. We wish to thank Dr. Douglas Moore for his expertise in micro-CT imaging and Geoffrey Williams for the use of the Leduc Bioimaging Facility at Brown University.

## Notes

### Competing Interest Statement

The authors have declared no competing interest.

### Summary of Updates

Added section on chronic in vivo experiments.

